# Increased NFAT and NFκB signalling contribute to the hyperinflammatory phenotype in response to *Aspergillus fumigatus* in cystic fibrosis

**DOI:** 10.1101/2024.11.29.625998

**Authors:** Amelia Bercusson, Thomas J Williams, Nicholas J Simmonds, Eric WFW Alton, Uta Griesenbach, Anand Shah, Adilia Warris, Darius Armstrong-James

**Author notes:** These authors contributed equally to this work (Amelia Bercusson, Thomas J Williams).

## Abstract

*Aspergillus fumigatus* (*Af*) is a major mould pathogen found ubiquitously in the air. It commonly infects the airways of people with cystic fibrosis (CF) leading to *Aspergillus* bronchitis or allergic bronchopulmonary aspergillosis. Resident alveolar macrophages and recruited neutrophils are important first lines of defence for clearance of *Af* in the lung. However, their contribution to the inflammatory phenotype in CF during *Af* infection is not well understood. Here, utilising CFTR deficient mice we describe a hyperinflammatory phenotype in both acute and allergic murine models of pulmonary aspergillosis. We show that during aspergillosis, CFTR deficiency leads to increased alveolar macrophage death and persistent inflammation of the airways in CF, accompanied by impaired fungal control. Utilising CFTR deficient murine cells and primary human CF cells we show that at a cellular level there is increased activation of NFκB and NFAT in response to *Af* which, as in *in vivo* models, is associated with increased cell death and reduced fungal control. Taken together, these studies indicate that CFTR deficiency promotes increased activation of inflammatory pathways, the induction of macrophage cell death and reduced fungal control contributing to the hyper-inflammatory of pulmonary aspergillosis phenotypes in CF.

**Author Summary:** The airways of people with cystic fibrosis (pwCF) are commonly colonised with *Aspergillus fumigatus* (*Af*) resulting in a persistent hyperinflammatory state and the development of allergy. Understanding how first line defence innate cells, such as alveolar macrophages and neutrophils, contribute to this hyperinflammatory phenotype is important in developing novel treatments to preserve lung function in pwCF. In this study, we report that CFTR deficiency leads to increased alveolar macrophage death and persistent inflammation of the airways in pwCF, which is associated with impaired control of infection. We identify the increased activation of the transcription factors NFκB and NFAT as the mechanism of increased inflammatory cytokine production in CFTR deficient cells. This work is the first step in describing molecular mechanisms of hyperinflammation in CF in response to fungal infection and lays the groundwork for further dissection of inflammatory pathways to target with immunotherapeutic approaches.

## Introduction

The major mould pathogen *Aspergillus fumigatus* (*Af*) is an air-borne saprophytic fungus that is found ubiquitously in the environment and continuously inhaled as spores (conidia) ^1^. Whilst healthy individuals are able to clear *Af* conidia, in individuals with chronic respiratory diseases or immunodeficiencies, acute or chronic forms of pulmonary aspergillosis may arise ^2–4^. Recent estimates suggest there are approximately 11.5 million people living with allergic bronchopulmonary aspergillosis (ABPA), 1.8 million with chronic pulmonary aspergillosis and more than 2 million with invasive pulmonary aspergillosis globally and with mortality rates of up to 18.5% and 85% for chronic and invasive pulmonary aspergillosis, respectively ^5^.

People with cystic fibrosis (pwCF), a disease caused by mutations in the cystic fibrosis (CF) transmembrane conductance regulator (CFTR) protein, are at increased risk of allergic bronchopulmonary aspergillosis, a chronic form of pulmonary aspergillosis characterised by Th2 responses in the airway and bronchiectasis ^6–10^. The CFTR protein regulates transepithelial ion flow of chloride and is required for the maintenance of the homeostatic volume of airway surface fluid ^11^. Loss of function of CFTR results in these ions being unable to cross the membrane, which leads to a thickened mucous layer causing lungs to lose their ability to maintain mucociliary clearance ^12,13^. Consequently, CF airways are characterised by the presence of chronic microbial contamination and inflammation ^14–17^. While there is no doubt that epithelial cell dysfunction contributes to this phenotype ^18–23^, there is increasing evidence that other components of the innate and adaptive immune responses are also involved in the pathogenesis of CF airways disease ^24–26,15,27–32^. At the airway epithelial surface, macrophages have a critical role in clearing pathogens and recruiting other immune cells during fungal infections. Alveolar macrophages, a group of lung resident cells, provide early defence in the airways against the conidia which are inhaled daily, efficiently controlling and clearing them ^33,34^. However, in instances of high fungal burden or fungal germination other immune cells are required to control the infection, the most prominent being neutrophils ^35–37^.

Microbe trapping by the thick mucus layer, characteristic of CF, results in an inflammatory neutrophil response, which leads to progressive lung disease and ultimately to pulmonary failure, the primary cause of death in CF ^38^. Evidence from bacterial studies in CF also show that CFTR dysfunction can lead to impaired microbe killing and greater inflammatory cytokine production in macrophages ^26,30–32^. This microbe trapping combined with impaired microbe killing is likely to lead to the airways of pwCF becoming colonised with filamentous fungi, of which *Aspergillus fumigatus* (*Af*) is most prevalent. Studies suggest that approximately 36% of pwCF have *A. fumigatus* colonisation of the airway, with between 30 to 65% being sensitised to *A. fumigatus*, determined by *A. fumigatus* specific IgE levels or skin prick tests ^6–10^.

In this study we aimed to evaluate the role of the CFTR protein in the inflammatory response to, and induction of cell death by *A. fumigatus* in both acute and chronic murine models of infection and cells from pwCF. Using both acute and chronic models of infection in CF mice we evaluated inflammatory cell recruitment and cytokine production in response to A. fumigatus. To accompany this in vivo data, we next examined how the regulation of inflammation in both murine and human macrophages during A. fumigatus infection was altered by CF. Finally, we investigated the potential for A. fumigatus to induce inflammatory macrophage cell in the context of CF and how this impacts fungal control in vitro.

## Methods

### Fungal Culture

*Aspergillus fumigatus* AF CEA10 (FGSC A1163) was obtained from the Fungal Genetics Stock Centre and was used for Western blot, ELISA, Luminex, fungal killing, LDH release and plate reader microscopy experiments. An eGFP-expressing strain (ATCC46645-eGFP, a gift from Frank Ebel) was used for confocal and widefield microscopy, flow cytometry and *in vivo* animal infections. All strains were cultured on Sabouraud dextrose agar (Oxoid). Conidia were harvested in 0.1% Tween/H_2_O and filtered through MIRACLOTH (Calbiochem, UK). Conidial suspensions were washed twice in sterile water and resuspended in RPMI 1640 at concentrations shown. To generate swollen conidia (SC), resting conidia (RC) were suspended in RPMI 1640 at 1.5 × 10^6^ conidia/ml and swollen at 37°C for 4 h.

### Animals

All mouse experiments were approved by the United Kingdom Home Office and the Imperial College London Animal Ethics Committee and performed in accordance with the project license PPL 70/7941. C57BL/6 male mice (6-8wks) were ordered from Charles River, UK. Wildtype (WT) and litter-mate homozygote knockout Cftr^tm1Unc^- Tg(FABPCFTR)1Jaw/J mice (CFTR^-/-^) (originally obtained from Jackson laboratories) were bred in house. These animals express CFTR under the control of the fatty acid binding promoter in the intestine and thereby do not manifest intestinal disease, which would otherwise prevent the longevity of these CF mice ^39^. All mice were housed in individually vented cages with free access to autoclaved food and water. WT and CFTR^-/-^ mice were co-housed with separation by gender and age cohort rather than genotype. For all experiments, mice were age- and-sex matched as closely as possible and were of similar starting weight pre-experiment.

### Bone marrow derived macrophage (BMDM) Generation

Wildtype (WT) C57BL/6 or CFTR knock-out mice (aged 8 to 10 weeks) were culled by overdose of phenobarbitone and the femur and tibia removed aseptically. Excess muscle was removed from the legs and the leg bones carefully cut open adjacent to each joint. A 28-G needle was inserted into the bone marrow cavity and the marrow flushed into a 50 ml Falcon tube using cold PBS without calcium or magnesium (PBS ^Ca-/Mg-^) supplemented with 2mM EDTA (Gibco by Life Technologies) until the bone appeared white. The cells were centrifuged for 10 min at 500g at room temperature and re-suspended in Red Blood cell lysis buffer (Sigma-Aldrich). After 10 min incubation, an equal amount of PBS was added, and the cells were spun down for 5 min at 500g. The remaining cell pellet was re-suspended in differentiation media consisting of RPMI 1640 supplemented with 10% FCS, 40 ng/ml recombinant murine M-CSF (eBioscience) and 200 IU/ml Penicillin/Streptomycin and counted by haemocytometer. 5 x 10^6^ cells were plated per 15 mm Petri dish in 8 ml of media and incubated at 37 C. After 3 days of culture, 4 ml of fresh differentiation media was added to each petri dish. After 7 days of culture, differentiated macrophages were washed once with PBS and harvested in PBS ^Ca-/Mg-^ supplemented with 2mM EDTA. For long-term storage, cells were re-suspended in RPMI 1640 supplemented with 10% FCS and 5% DMSO, frozen at −80^0^C overnight and stored in the vapour phase of liquid nitrogen. For culture after differentiation, macrophages were thawed and re-suspended in culture media consisting of RPMI containing 10% FCS, 20 ng/ml recombinant murine M-CSF and 200 IU/ml Penicillin/Streptomycin, seeded and rested for 2 days before being used for experimentation. On the day before experimentation, the medium was replaced with culture media supplemented by 200 IU/ml IFN-γ for 24 hours. On the day of experimentation, the IFN-γ was removed.

### Human monocyte derived macrophage (hMDM) Generation

Collection and study of human macrophages was approved by the Biomedical Research Unit Biobank research project (NRES reference 10/H0504/9), Royal Brompton and Harefield NHS Trust (AS1). All experiments conformed to the principles set out in the WMA declaration of Helsinki and the Department of Health and Human Services Belmont Report.

Peripheral blood mononuclear cells (PBMCs) were isolated from blood from healthy volunteers or pwCF mixed 3:1 in warm RPMI and layered over Ficoll-Paque PLUS (GE healthcare). Samples were centrifuged at 450g for 45 minutes and the PBMC layer removed. PBMCs were incubated with red cell lysing buffer Hybri-Max (Sigma) for 10 minutes and then washed at 200g in warm PBS x 2 to remove platelets. Monocytes were subsequently isolated by negative magnetic bead selection using a pan-monocyte isolation kit (Miltenyi Biotech, Auburn, CA). To obtain monocyte-derived macrophages (MDMs), freshly isolated monocytes were differentiated in RPMI 1640 medium supplemented with 10% human serum (Sigma, UK), 200 IU/ml Penicillin-Streptomycin and 5 ng/ml granulocyte macrophage colony-stimulating factor (GM-CSF) (Peprotech, UK) for 7 days at 37°C.

### Wide-field Fluorescent Microscopy

Cells were seeded on glass coverslips in 24-well plates and treated as indicated. Before staining, cells were washed in PBS x1 and fixed in 2% PFA for 15 minutes followed by quenching in 50mM NH_4_Cl for 10 min. Cells were blocked and permeabilised in PBS containing 10% goat serum (Sigma-Aldrich) and 0.1% Saponin (Sigma-Aldrich) for 2h at room temperature and incubated overnight at 4°C with a primary antibody diluted in the blocking and permeabilization buffer (see table). After washing with PBS three times, cells were incubated with anti-rabbit Cy5 or anti-mouse Cy5 antibodies (Life Technologies) for 45min at room temperature in the dark, mounted with Vectashield mounting medium containing DAPI (Vector laboratories) and sealed with nail varnish to prevent evaporation. Image analysis was performed using Image J software. NFAT and NFκB translocation studies were conducted with a Zeiss LSM-510 widefield microscope. Nuclear translocation was quantified by determining the percentage overlap of the NFAT/NFκB channel with the DAPI channel (the Mander’s co-efficient) using a macro written for the project (thanks to Stephen Rothery, Imperial Facility for Imaging by Light Microscopy).

### LDH Assay

LDH levels in tissue culture supernatants were measured by colorimetric formazan product formation using the CytoTox 96® Non-Radioactive Cytotoxicity Assay kit (Promega) according to the manufacturer’s instructions.

### Plate Reader Fluorescent Microscopy (Fluorescent Apoptosis/Necrosis Assay (FAN Assay)

Macrophages were seeded in in black-walled clear-bottomed 96-well microplates (Corning). On day of experiment, *A. fumigatus* conidia were added to appropriate wells and plates were centrifuged at 100 g for 3 min. Plates were incubated at 37° C for 1 hour. RPMI containing Propidium iodide (PI) and Ac-DEVD-AMC (Enzo Life Sciences) was added to each well and PI-positive cell death and AMC cleavage by Caspase 3 (an analogue measure of apoptosis) were measured over 16 hrs by time-lapse fluorescence microscopy using a TECAN Infinite PRO microplate reader.

### ELISA and Luminex

For the quantification of cytokines in tissue culture and BAL supernatants, ELISAs were performed according to the manufacturer’s instruction using the TNF-α, CXCL1 and IL-1β DuoSet kits from R& D systems. In addition, cytokines in supernatants were measured with a Meso Scale Discovery (MSD) multiplex assay kit Containing antibodies for the following analytes: IFN-γ, IL-6, TNF-α, MIP-2, MCP-1, IL-1β, CXCL1, CXCL10, IL-5, IL-33.

### CFU

For *in vitro* killing assays, cells were infected with *A. fumigatus* swollen conidia (MOI=1) for 6 hrs. Cells were then washed to remove extra-cellular conidia and lysed in 0.1% Tween 20/dH2O. Serial dilutions were plated onto Sabouraud agar plates. Plates were incubated at 37° C overnight and CFUs were counted the next day. For fungal burden in BAL samples, serial dilution of BAL fluid was performed in 0.1% Tween 20/dH2O and plated onto Sabouraud agar plates. Plates were incubated at 37° C overnight and CFUs were counted the next day.

### Flow Cytometry

#### Phagocytosis

Cells were seeded in 24-well plates and on the day of experimentation infected with biotinylated e-GFP *A. fumigatus* conidia at an MOI of 1 for 2 hours. Unattached conidia were washed off and cells fixed in 2% PFA for 15 minutes followed by quenching in 50mM NH_4_Cl for 10 min. Cells were counterstained with Cy3 biotin antibody. Flow Cytometry was performed on an LSR Fortessa flow cytometer (BD Biosciences). Data was analysed using FlowJo software (Oregon, USA).

#### Cell Influx

Murine bronchoalveolar lavage fluid was spun down at 500 g for 10 min and the cell pellet resuspended in PBS containing an Fc receptor blocking antibody (anti-CD16/CD32, clone 93, eBio-science, 1:100). Single-cell suspensions were stained with Live/Dead Fixable Blue Stain (L34959, Thermo Fisher Scientific) and combinations of the following surface anti-mouse markers: CD19-BV785 (1:100, 115543, Biolegend), CD45.2- BUV737 (1:400, 564880, BD), CD3-BUV395 (1:200, 740268. BD), Ly6C-PE-Cy7 (1:200, 25-5932-82, eBioscience), MHC-II-APC-Cy7 (1:200, 107628, Biolegend), Ly6G-AF700 (1:200, 127622, Biolegend), CD11b-BV605 (1:200, 101237, Biolegend), F4/80-APC (1:200, 17-4801-82, Invitrogen), CD11c-FITC (1:200, 557400, BD), CD11c-PE-Cy7 (1:200, 558079, BD), Siglec F-PE (1:200, 562068, BD) and fixed in 2% PFA. Flow Cytometry was performed on an LSR Fortessa flow cytometer (BD Biosciences). Data was analysed using FlowJo software. The gating strategy for identifying lung immune cells by flow cytometry is given in Supplemental Methods Figure 1.

**Figure 1:**
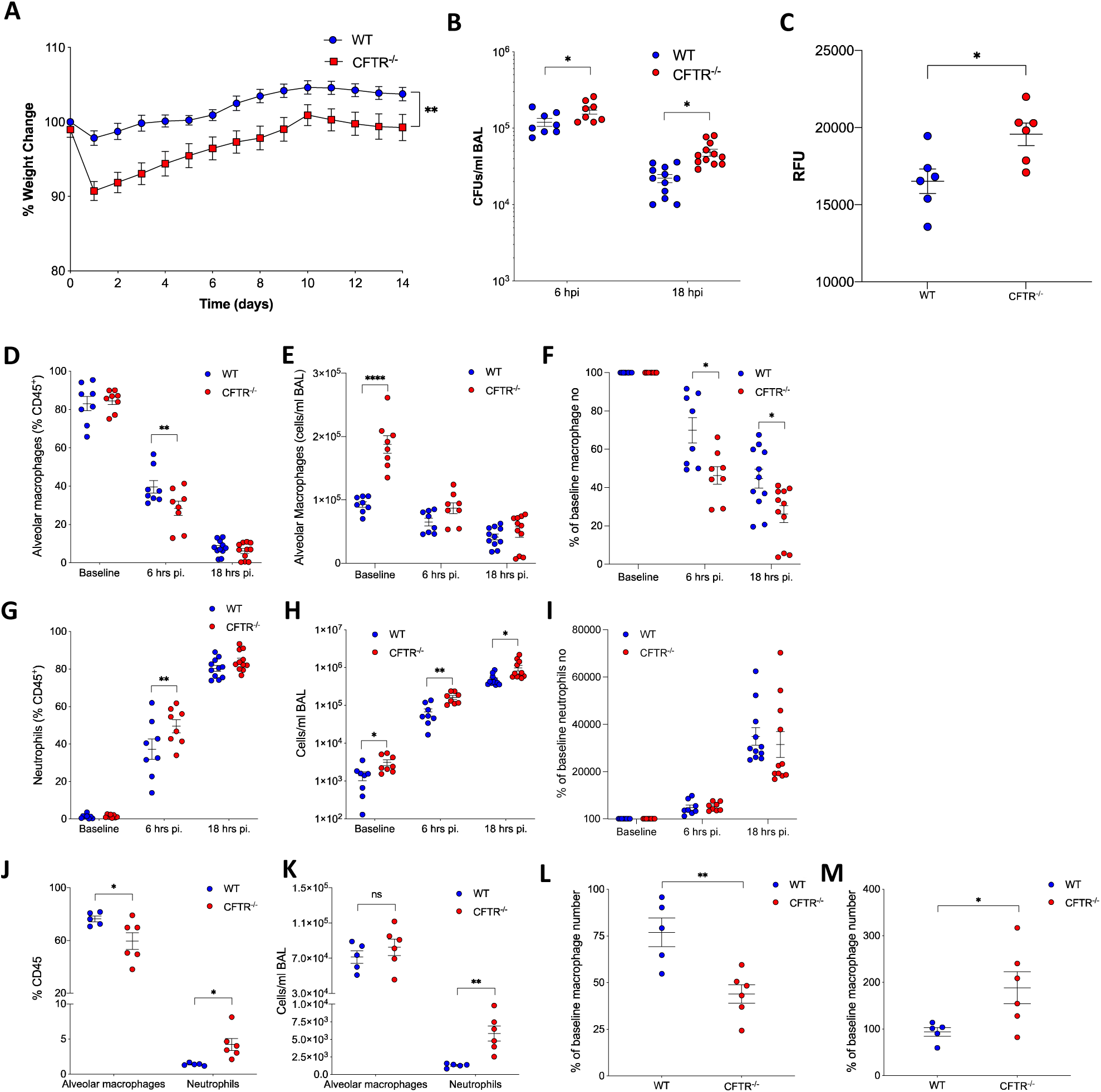
(**A**) WT and CFTR^-/-^ mice were infected intranasally with 1 x 10^7^ eGFP *A. fumigatus* (**A**) body weight was monitored daily for 3 weeks post infection (n=8). (**B-I**) WT and CFTR^-/-^ mice were infected intranasally with 1 x 10^7^ eGFP *A. fumigatus* conidia and sacrificed at 6 and 18 hours post infection for bronchoalveolar lavage. (**B**) Colony forming units (CFU) were determined in BAL fluid at the time points indicated. (**C**) GFP^+^ alveolar macrophage population was identified by flow cytometry and geometric mean of GFP fluorescent intensity was calculated (n=8-12). (**D-I)** Cell influx was studied by flow cytometry, assessing (**D-F**) macrophages and (**G-I**) neutrophils as (**D, G**) proportion of CD45^+^ cells, (**E, H**) absolute count of cells/ml BAL fluid and (**F, I**) percentage of average baseline numbers (n=8-11). (**J-M**) WT and CFTR-/- mice were infected with 1×10^7^ resting conidia and sacrificed 3 weeks post-infection. Cell influx was studied by FACS, assessing (**J**) alveolar macrophages and neutrophil proportions as % of CD45^+^ cells. (**K**) Absolute count of alveolar macrophages and neutrophils/ml BAL fluid. (**L**) Percentage of average baseline macrophage proportions (**M**) % of average baseline neutrophil numbers (n=8- 11). Data are represented as mean ± SEM. p values calculated by two-way ANOVA (**A**) Students T test (**B, C, L, M**) and one-way ANOVA (**D-K**), *p<0.05, **p<0.01, ***p<0.001, ****p<0.0001, ns – non-significant. BAL – Bronchoalveolar Lavage, WT – Wild Type, pi – post-infection.

### Infections

#### Acute model

Mice were infected with 1×10^7^ of either CEA10 or e-GFP *A. fumigatus* live resting conidia administered intranasally under isoflurane anaesthesia in 50 μl of PBS. Mice were culled at 6 hours, 18 hours or 3 weeks post infection by intra-peritoneal overdose of pentobarbital.

#### Allergic Model

Mice were infected with 2 x 10^5^ live resting conidia of *Af* CEA10 administered intranasally in 50 µl of PBS while under isoflurane anaesthesia. Doses were repeated daily for 14 days. Mice were culled 24 hours after the final dose by intra-peritoneal overdose of pentobarbital.

#### Bronchoalveolar lavage

Mice were culled prior to bronchoalveolar lavage by intra-peritoneal overdose of phenobarbitone. Bronchoalveolar lavage (BAL) was performed by tracheal cut-down and intubation with a self-designed intubation catheter. Lungs were lavaged by installing 3 x 800 μl of PBS/2mM EDTA. The lavage fluid was stored on ice until further processing. 110 μl was removed for fungal burden analysis by CFU assay (see above). Cells and supernatant were separated by spinning at 400 g for 10 min. Cell pellets were processed immediately for flow cytometry while supernatants were stored at −80 C for further analysis.

#### Quantification and statistical analysis

All data are expressed as mean ± SEM. Statistics were calculated using Prism software (version 7.0; GraphPad) using one-way ANOVA (comparison of ≥ 3 groups), two-way ANOVA and Student’s t test (comparison between 2 groups) were carried out. For all figures, p values are represented as followed: ∗p < 0.05; ∗∗p < 0.01; ∗∗∗p < 0.001, ∗∗∗∗p < 0.0001

## Results

### Acute Aspergillus infection leads to weight loss, alveolar macrophage loss and increased neutrophil recruitment in cystic fibrosis

PwCF exhibit hyperinflammatory cytokine responses and inflammatory cell recruitment in response to infection ^26,31,30,32^. To investigate the impact of CF on acute immune responses to *Af*, we exploited the Cftr^tm1Unc^-Tg(FABPCFTR)1Jaw/J (CFTR^-/-^) mice to model the acute airway inflammatory response to *Af* challenge.

CFTR^-/-^ and WT control mice were infected intranasally with 1 x 10^7^ eGFP *A. fumigatus* (ATCC46645-eGFP) resting conidia and monitored over a three-week period. CFTR-/- mice lost significantly more weight than their WT counterparts and were slower to recover to their baseline weight (**Fig 1A**, p≤0.05). CFUs in bronchoalveolar lavage fluid from infected CFTR-/- and WT mice, obtained 6- and 18- hours post-infection, were assessed. Fungal survival was significantly increased in CFTR^-/-^ mice compared to controls at both time-points (**Fig 1B**, p≤0.05). The geometric mean fluorescent intensity of the eGFP-*A. fumigatus* in alveolar macrophage cells was also significantly increased in CFTR^-/-^ compared to WT mice, identifying an alveolar macrophage-specific killing defect *in vivo* (**Fig 1C**, p≤0.05). Flow-cytometry based cellular analysis of BAL fluid after 6 and 18 hours post infection demonstrated that although alveolar macrophage numbers decreased in both groups post-infection, there was a significantly greater loss of cells from baseline in the CFTR-/- mice than in WT, suggesting increased *Aspergillus*-induced macrophage cell death (**Fig 1D-F**, p≤0.05). CF is recognised as a neutrophilic airways disease and consistent with this we found that neutrophil recruitment to the airways was significantly increased in CFTR^-/-^ mice compared to wild type (WT) controls at both 6- and 18-hours post-infection (**Fig 1G-I**, p≤0.05).

To assess whether resolution of the inflammatory cell changes induced by *Aspergillus* infection is delayed in CF, WT and CFTR^-/-^, mice were examined at 3 weeks post-infection. Flow cytometric analysis of bronchoalveolar lavage fluid showed an increased proportion and absolute number of neutrophils in the airways at 3 weeks post infection in CFTR^-/-^ compared to WT controls (**Fig 1J, K**, p≤0.05) Alveolar macrophages were significantly delayed in returning to baseline numbers (**Fig 1L**, p≤0.05), whereas neutrophils were still significantly increased in CFTR^-/-^ airways at three weeks compared to controls where they returned to baseline numbers (**Fig 1M**, p≤0.05). These data shows that there are significant changes in the immune profile of the lung in response to *A. fumigatus* infection in CFTR^-/-^ mice compared to WT controls early in infection and this altered response persists up to 3 weeks.

### Acute Aspergillus infection leads to an increased inflammatory profile of the airways in cystic fibrosis knock-out mice

Critical to the recruitment of neutrophils to the airways is the early release of chemotactic cytokines such as IL-8/CXCL1, TNF-α and MIP-2 and other inflammatory cytokines. To further interrogate the changes in inflammatory response in CFTR-/- mice, Meso Scale Discovery (MSD) multiplex analysis was undertaken on bronchoalveolar lavage samples from mice intranasally infected with 1 x 10^7^ *A. fumigatus* resting conidia and sacrificed after 6 and 18 hours post infection. This revealed increased CXCL1 release at 6 hours post-infection in CFTR^-/-^ mice compared to controls. MCP-1 and CXCL10 are classically know as monocyte chemokines; however, there is evidence from murine infections studies that they also contribute to neutrophil chemotaxis and were also found to be significantly increased in CFTR-/- samples compared to WT controls. Increased release of inflammatory cytokines was also observed at 18 hours post-infection with MSD analysis revealing raised levels of MIP-1a, MIP-2, MCP-1, TNF-α, IL-1β, IL-6 and IL-33 in BAL samples from CFTR^-/-^ mice compared to controls (**Fig 2A-J**, p≤0.05). Overall, these observations indicate that the secretion of chemokines required for inflammatory cell recruitment is increased early in infection in BAL from CFTR^-/-^ mice compared to controls and that the presence of fungus in the airways of CFTR^-/-^ mice leads to greater inflammatory cytokine production, allowing for ongoing immune stimulation.

**Figure 2:**
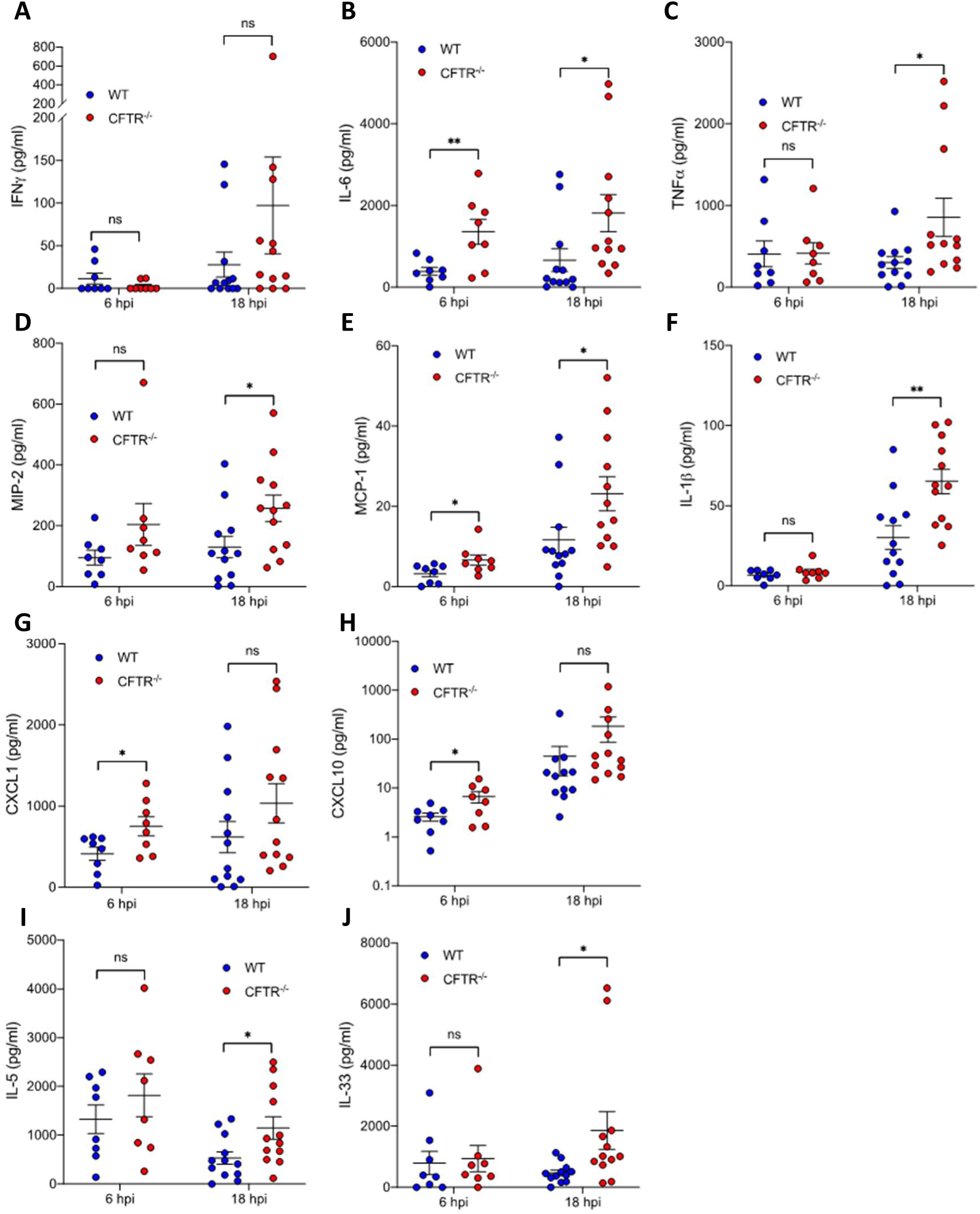
(**A-J**) WT and CFTR^-/-^ mice were infected intranasally with 1 x 10^7^ eGFP *A. fumigatus* conidia. Mice were culled at 6 (n=8) and 18 hours (n=12) and bronchoalveolar lavage (BAL) was performed. Cytokine levels in the BAL fluid were measured by MSD for (**A**) IFN-y, (**B**) IL-6, (**C**) TNFa, (**D**) MIP-2, (**E**) MCP-1, (**F**) IL-1b, (**G**) CXCL1, (**H**) CXCL10, (**I**) IL-5 and (**J**) IL-33. Data represented as mean ± SEM. p values calculated by Students T test, *p<0.05, **p<0.01, ns – non-significant. BAL – Bronchoalveolar Lavage, , MSD – meso scale discovery, WT – Wild Type; hpi – hours post-infection.

### CFTR^-/-^ mice exhibit increased weight loss, alveolar macrophage death and increased neutrophil recruitment during allergic pulmonary aspergillosis

ABPA is one of the most prevalent forms of aspergillosis in CF, with a prevalence up to 15% ^40,7,41,42^. Therefore, we next aimed to investigate whether the hyperinflammatory phenotype seen in acute *A. fumigatus* infection was also observed during allergic pulmonary aspergillosis. We employed a repeat challenge model of allergic aspergillosis in which WT and CFTR^-/-^ mice were intranasally infected with 2 x 10^5^ CEA10 *A. fumigatus* conidia daily for 14 days. No mortality was observed in either the WT or CFTR^-/-^ groups that were infected with *A. fumigatus*. However, CFTR^-/-^ mice lost a significant amount of weight compared to both the PBS CFTR^-/-^ controls and the WT *A. fumigatus* infected mice along the time course of the experiment, whereas there was no significant weight loss seen between WT PBS treated and *A. fumigatus* infected groups (**Fig 3A**, p≤0.01). In both WT and CFTR^-/-^ mice the temporal changes in BAL fluid between repeatedly infected and PBS-treated mice was similar with increased levels of eosinophils, neutrophils, and T cells, while alveolar macrophages were decreased (**Fig 3B-E**). While there were no significant differences in the proportion or number of eosinophils or the loss of alveolar macrophages between WT and CFTR^-/-^ mice (**Fig 3B, C**), it was observed that CFTR^-/-^ mice repeatedly infected with *A. fumigatus* had both a higher proportion and larger population of recruited neutrophils (**Fig 3D**, p≤0.05) and T cells (**Fig 3E** p≤0.0001) in the airways. These data suggests that in a model of allergic aspergillosis, CFTR^-/-^ mice have greater recruitment of cells to the airways that may contribute to a hyperinflammatory phenotype compared to WT controls.

**Figure 3:**
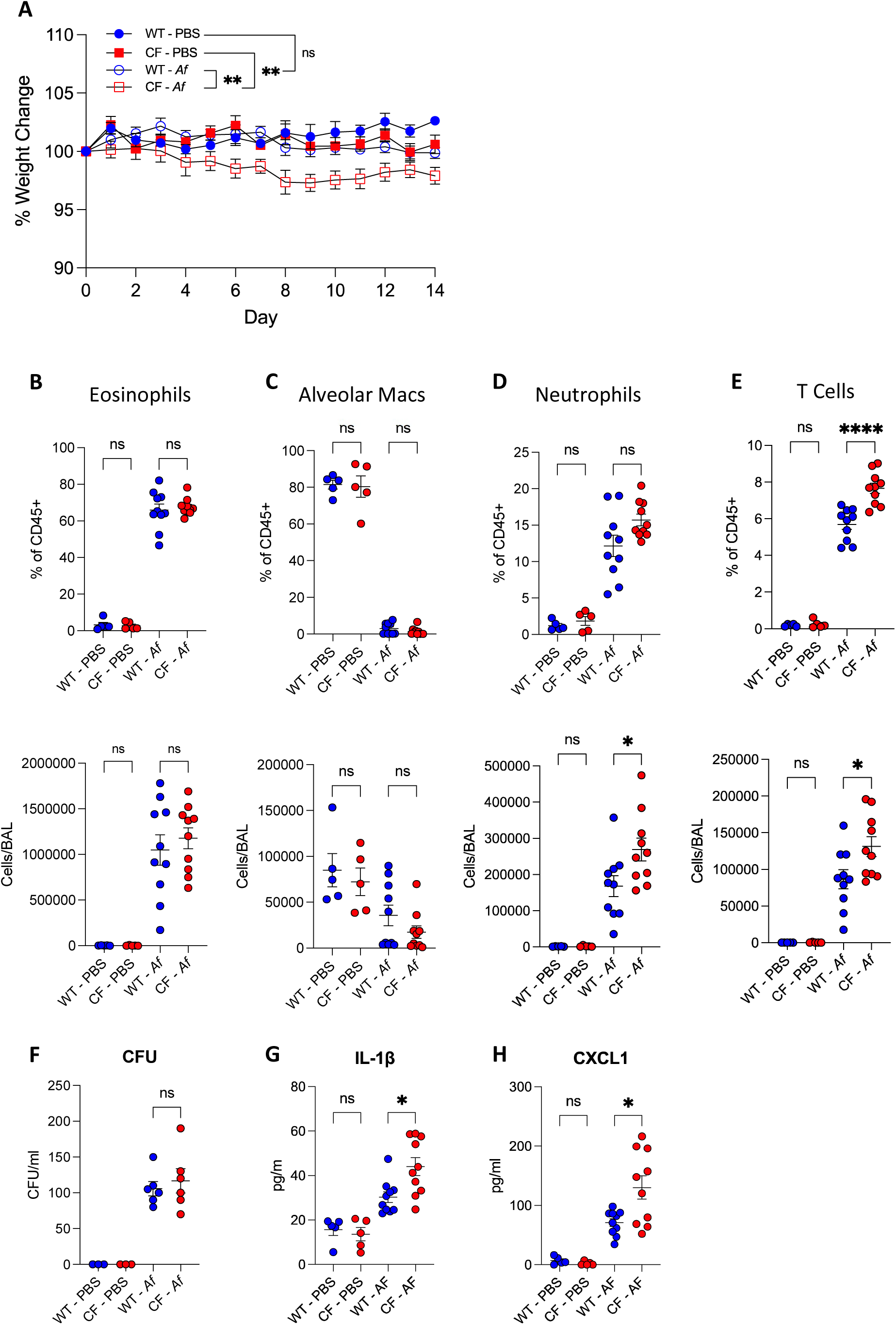
(**A-H**) WT or CFTR-/- mice were dosed daily with 2×10^5^ CEA10 *A. fumigatus* or PBS for 14 days and assessed for (**A**) weight change (n=5-10). Mice were culled 24 hours after the final does and BAL was carried out. Flow cytometry was utilised to determine the immune cell populations in the airways. Percentage and enumeration of cells for (**B**) Eosinophils, (**C**) Alveolar Macrophages, (**D**) Neutrophils and (**E**) T Cells (n=5-10). BAL fluid was assessed for (**F**) CFUs and the inflammatory cytokines (**G**) IL-1β and (**H**) CXCL1 were assessed by ELISA (n=5-10). Data represented as mean ± SEM. (**A**) Two-way ANOVA(**B-H**) One-way ANOVA, *P<0.05, **P<0.01, ***P<0.001, ****P<0.0001, ns – not significant. BAL – Bronchoalveolar Lavage, WT – Wild Type.

CFU assays were carried on the BALF to investigate whether these changes in immune cell recruitment impacted fungal control. No significant differences in CFUs were observed between repeatedly exposed WT and CFTR^-/-^ mice (**Fig 3F**).

Next IL-1β, a traditional marker of inflammation and pore forming cell death, and CXCL1, the neutrophil chemoattractant were measured by ELISAs on the BALF supernatant to examine whether the increased inflammatory immune cell recruitment was accompanied by increase inflammatory cytokine release. CFTR^-/-^ mice were observed to have significantly more IL-1β and CXLC1 in the BALF compared to the WT controls (**Fig 3G, H**, p≤0.05). Taken together these data suggest that in allergic pulmonary aspergillosis CFTR^-/-^ mice have increased release of IL-1β and CXCL1, resulting in the increased recruitment of neutrophils to the airway. However, this excessive cytokine release does not result in increased fungal clearance thereby creating a hyperinflammatory environment with a limited effect on controlling infection.

### The hyperinflammatory phenotype seen in CF is driven by increased NFκb and NFAT signaling in the absence of CFTR

Upon encountering macrophages within the airway, *A. fumigatus* conidia are rapidly taken up into endosomes. As the endosome matures, several signalling pathways, including NFAT and NF-κB, are responsible for coordinating the cell’s response to the fungus ^43^. We have previously shown that calcineurin-NFAT pathway plays a key role in regulating cytokine release, fungal killing, kinetics of phagosomal maturation and programmed cell death pathways ^44,45^. As we observed increased inflammatory cytokine production by CFTR^-/-^ macrophages, we next investigated whether this was related to altered inflammatory transcription factor responses. We found that both NFAT and NF- κB activation were significantly increased in CFTR^-/-^ bone-marrow derived macrophages (BMDM) compared to WT controls in response to *A. fumigatus* infection, as assessed by nuclear colocalization of the transcription factor (**Fig 4A-D**, p≤0.05). Both NFAT and NFκB are involved in regulating cytokine and chemokine secretion by macrophages in response to *A. fumigatus*. Consistent with our *in vivo* observations, increased activation of NFAT and NF-κB in CFTR^-/-^ macrophages correlated with increased levels of *Aspergillus*-induced inflammatory cytokines, including TNF-α, CXCL1 and IL-1β, detected in the cell culture supernatants of CFTR^-/-^ BMDMs compared to WT controls (**Fig 4E**, p≤0.05). These findings were confirmed in human monocyte-derived macrophages from pwCF which showed a similar increase in activation of NFAT and NF-κB in response to *A. fumigatus* infection compared to healthy controls, again shown by colocalization of the transcription factor stains (**Fig 4F**, p≤0.05). These data demonstrate there is increased activation of both NFAT and NF-κB in response to *A. fumigatus* in macrophages from CFTR^-/-^ mice as well as from pwCF *in vitro*.

**Figure 4:**
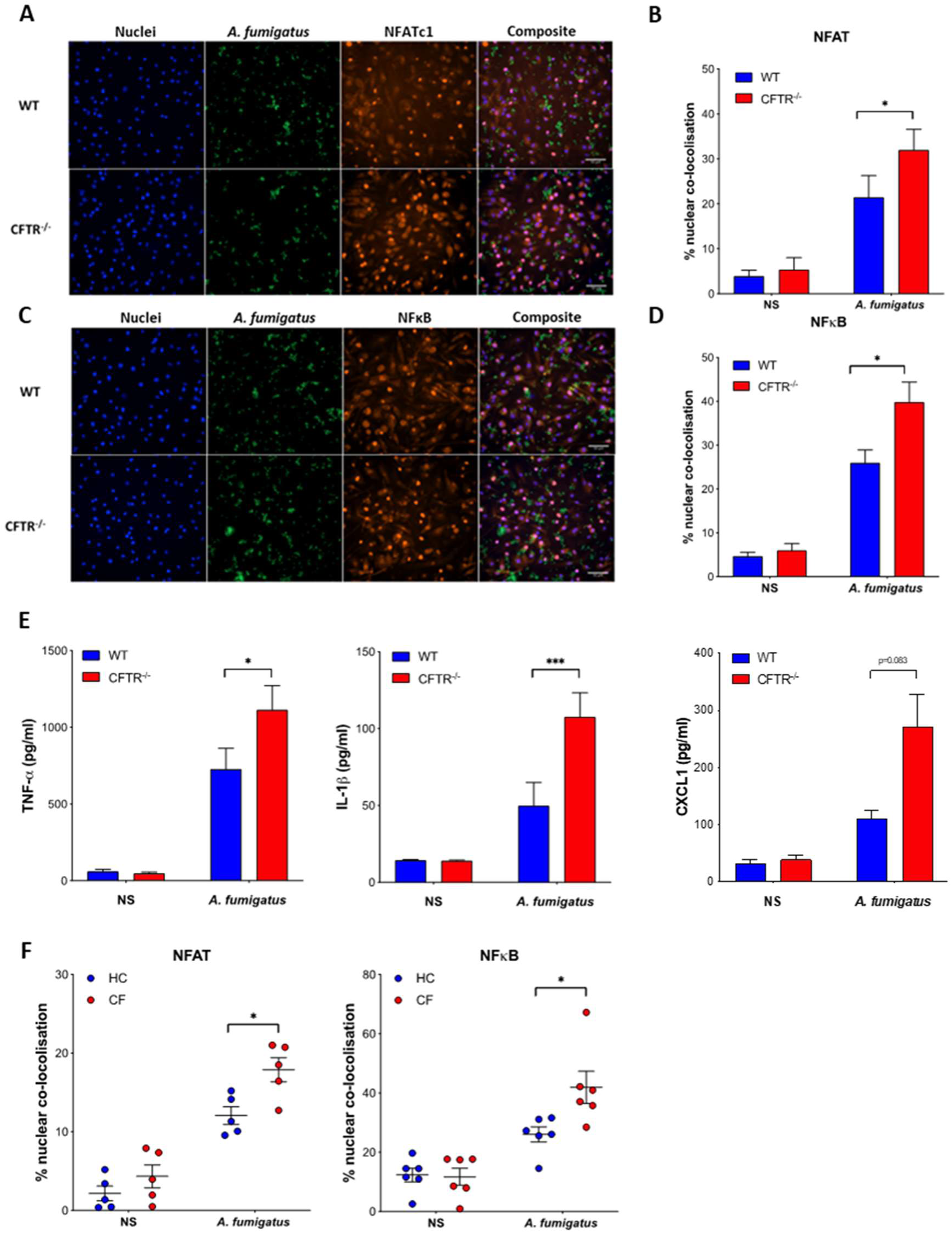
(**A-D**) WT and CFTR^-/-^ bone-marrow derived macrophages (BMDM) were stimulated with eGFP *A. fumigatus* (green) swollen conidia (MOI=1) for 1 hour and stained for NFAT or NFκB (orange) and DAPI nuclear stain (blue). Nuclear translocation was quantified by calculating the % overlap of the nuclear and transcription factor-conjugated fluorophore channels. Representative microscopy and quantification of translocation of (**A-B**) NFAT and (**C-D**) NFκB. Data were collected from at least 4 fields of view taken at random per condition per experiment (n=3). (**E**) WT and CFTR^-/-^ BMDMs were stimulated with *A. fumigatus* swollen conidia (MOI=1) for 16 hrs. The levels of secreted TNF-α (n=4), IL-1β (n=3) and CXCL1 (n=3) in cell culture supernatants were measured by ELISA. (**F**) Monocyte-derived macrophages from CF patients and healthy controls (HC) were stimulated with eGFP *A. fumigatus* swollen conidia (MOI=1) for 1 hour and transcription factor translocation was quantified for NFAT (n=5) and NF-κB (n=6) as in (**A-D**). Data represented as mean ± SEM. p values calculated by two-way ANOVA *p<0.05, **p<0.01, ***p<0.001. CF – Cystic Fibrosis, HC – Healthy Control, MOI - Multiplicity of Infection, NS – non-stimulated, WT – Wild Type.

### The hyperinflammatory phenotype of CFTR loss is accompanied by increased inflammatory cell death and reduction of fungal control by macrophages

Upon infection cells may undergo some form of programmed cell death, which can be broadly divided into non-inflammatory cell death, a prime example being apoptosis, and inflammatory cell death, such as necroptosis and pyroptosis ^46^. To investigate if cell death outcomes were altered by loss of CFTR, BMDMs from CFTR^-/-^ and WT mice were infected with *A. fumigatus* conidia *in vitro* and cell death was assessed using a time-lapse fluorescent plate reader with propidium iodide (PI), a nucleic acid stain, as a marker of necrosis and Ac-DEVD-AMC, a fluorogenic substrate of Caspase-3, as a marker of apoptosis. PI^+^ necrotic cell death was increased in CFTR^-/-^ macrophages compared to WT controls at 16 hours, but no difference was observed in apoptotic death (**Fig 5A-B**, p≤0.01). This increase in total death was confirmed by investigating lactate dehydrogenase (LDH) release; increased levels of cell death were observed in CFTR-/- BMDMs compared to controls (**Fig 5C**, p≤0.05). Cells that do not kill an invading pathogen, can undergo ‘programmed cell death’ in an attempt to prevent the pathogen escaping and damaging surrounding cells ^46^. To investigate whether there was a link between increased cell death and fungal control, phagocytosis and fungal killing were assessed. To examine whether there was an intrinsic CFTR-related defect in phagocytosis, CFTR ^-/-^ and WT BMDMs were incubated with biotinylated eGFP *A. fumigatus* for 2 hours then counterstained with an anti-biotin Cy3 antibody and phagocytosis was measured by flow cytometry, with phagocytosed conidia determined as GFP^+^Cy3^-^ cells. We did not observe any significant difference in phagocytosis between the WT and CFTR ^-/-^ BMDMs (**Fig 5D**). Although uptake of fungal conidia was not impaired in CFTR^-/-^ macrophages, we found that fungal killing was impaired at 6 hours post-infection. The number of viable conidia extracted from CFTR^-/-^ macrophage cultures, assessed by CFU assays, was increased compared to those from WT macrophages, suggesting a killing defect (**Fig 5E**, p≤0.05). This was also observed in time-lapse microscopy using eGFP *A. fumigatus*, where greater fungal outgrowth was observed when CFTR^-/-^ macrophages were infected compared to WT controls (**Fig 5F**, p≤0.05). Therefore, these data show that CFTR^-/-^ macrophages exhibit increased cell death in response to *A. fumigatus* and an impaired ability to control the fungus *in vitro*.

**Figure 5:**
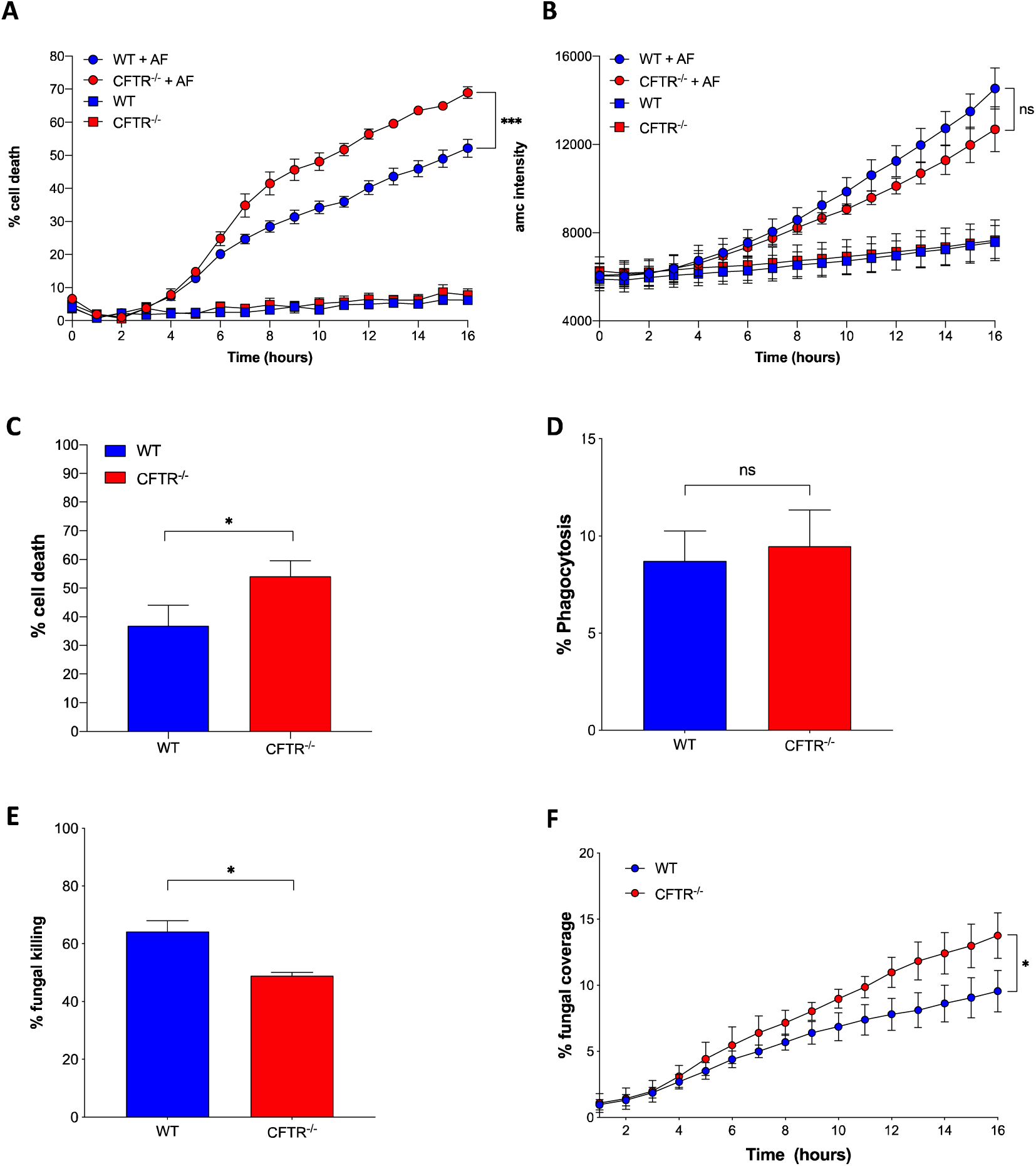
(**A + B**) BMDMs from WT and CFTR^-/-^ mice were infected with eGFP *A. fumigatus* swollen conidia (MOI=1) and PI (necrotic cell death) and Ac-DEVD-AMC Caspase-3 Substrate (apoptotic cell death) were added to the media. Plates were incubated over 16 hours in a fluorescence plate reader. (A) Percentage cell death quantified by comparison to 100% death control. (B) Levels of apoptosis were quantified by Ac-DEVD-AMC intensity. (**C**) BMDM from WT and CFTR^-/-^ mice were infected with eGFP *A. fumigatus* swollen conidia (MOI=1) for 18 hours. Percentage cell death was quantified by LDH levels in cell culture supernatants compared to a maximum release control. (**D**) WT and CFTR ^-/-^ BMDM were infected with biotinylated GFP *A. fumigatus* swollen conidia (MOI=1) for 2 hours. Cells were then counterstained with an anti-biotin Cy3 antibody and phagocytosis was measured by flow cytometry. (**E**) WT and CFTR ^-/-^ BMDMs were infected with eGFP *A. fumigatus* swollen conidia (MOI=1) for 6 hours. Percentage fungal killing was determined by CFU assay. (**F**) BMDMs from WT and CFTR^-/-^ mice were infected with eGFP *A. fumigatus* swollen conidia (MOI=1) and live time-lapse imaging was carried out over 16 hours. The % coverage of the field of view by GFP expressing *A. fumigatus* was measured. Data represented as mean ± SEM. p values calculated by paired Student t test (**C-E**), two-way ANOVA (**A, B, F**) *p<0.05, **p<0.01, ***p<0.001. CFU – Colony forming unit, MOI - Multiplicity of Infection, WT – Wild Type.

## Discussion

CFTR channel dysfunction likely results in excessive inflammatory responses to infection often leading to progressive bronchiectasis and decline in lung function in pwCF ^38^. Chronic infection and allergic inflammation with the major fungal pathogen *Aspergillus fumigatus* are major complications in pwCF leading to accelerated lung function decline and morbidity ^47–49^. In this study we utilised a combination of *in vivo* models of pulmonary aspergillosis alongside primary human cells and isolated murine macrophages to characterise innate immune responses to *A. fumigatus* in CF and their contribution to the well-recognized hyperinflammatory phenotype seen clinically in pwCF. In this study, we identify CF associated increased activation of pro-inflammatory pathways and release of pro-inflammatory cytokines in response to infection with *A. fumigatus*. Furthermore, we show that CFTR deficiency promotes the induction of macrophage cell death and reduction of fungal control.

First, we characterised the inflammatory response in both an acute and allergic model of aspergillosis to understand how different levels of fungal exposure may influence the inflammatory response in CF. In both the acute and allergic models of aspergillosis, CFTR^-/-^ mice lost more weight than the WT control group suggesting that infection with *A. fumigatus* induced a more potent and prolonged inflammatory response in the CFTR^-/-^ mice compared to wild type. In the acute model, weight loss was immediate and prolonged compared to the allergic model in which weight loss started to occur markedly after a week of repeated infections. This suggests that *A. fumigatus* is able to induce sustained inflammation in CF even in the absence of ABPA. Whilst only the acute model of aspergillosis showed marked loss of alveolar macrophages, both acute and allergic models exhibited increased influx of neutrophils to the infected airway in CFTR^-/-^ mice compared to WT controls, matching the neutrophil influx phenotype commonly associated with CF ^50^. Despite this increase in neutrophil numbers in the acute model, fungal control was reduced in CFTR^-/-^ compared to WT controls. Previous studies *in vitro* and *in vivo* have highlighted reduced microbial control in the absence of CFTR through impairment of microbial killing by macrophages, additionally it has been reported that CF epithelial cells infected with *A. fumigatus* have reduced fungal control ^31,51–53^. Furthermore, aberrant killing of *A. fumigatus* has also been demonstrated in COPD derived epithelial cells ^54^, showing the importance of understanding not only the innate immune response but also the initial epithelial response and how this contributes to the control of infection in these models. In line with previous CF infection models, both models of aspergillosis showed increased and sustained levels of inflammatory cytokines in the CFTR^-/-^ mice compared to WT controls, again highlighting the potent inflammation that can be induced by a single exposure to *A. fumigatus* in CF without an ABPA phenotype^55,56^.

To better understand the underlying basis for these observations, we further examined the observed *in vivo* increase in alveolar macrophage loss and cytokine production seen in CF *in vitro*. We found that *A. fumigatus* triggered the activation of the key transcription factors NFAT and NFκB in a significantly greater proportion of infected CF macrophages than wild-type macrophages. These findings were confirmed in human as well as murine macrophages. Whilst increased NFκB hyper-reactivity has been previously reported in CF epithelial cells and in CF macrophages to bacterial stimuli, to our knowledge this is the first description for a fungal pathogen and the first report of increased NFAT activation in CF macrophages in response to an infective stimulus ^57–59^. Notably, NFAT activation occurs in response to sustained calcium influx into the cytoplasm ^60^. Calcium signalling has been found to be dysregulated in CF epithelial cells with several mechanisms being identified as an explanation ^61–63^. Regulation of NFκB activation is complex and influenced by several different signalling pathways including calcium flux ^64^. Consistent with our observations, CFTR has been described as a negative regular of NFκB-mediated inflammatory signalling with cell membrane localisation of a functional CFTR ion channel necessary to prevent hyper-activation of NFκB activation pathways ^65^. One of the ways wild-type CFTR influences NFκB is via promoting the degradation of TRADD (TNF receptor 1 - associated death domain), which forms part of a multi-protein complex that enables the phosphorylation of the inhibitory protein IκBα ^18^.

The combination of increased NFAT and NFκB activation resulted in increased release of pro-inflammatory cytokines including TNF-α, CXCL1 (one of the murine equivalents of human IL-8), and IL-1β. Both TNF-α and CXCL1 are potent neutrophil chemokines, providing a likely explanation for the neutrophilic airways disease observed in our models. IL-1β is released following inflammasome activation and increased NLRP3 inflammasome activity has been noted in murine and human CF cells ^55^. These *in vitro* findings using infection-naïve macrophages support the hypothesis that intrinsic CFTR-related abnormalities in macrophage signalling pathways contribute to the generation of inflammation in CF airways.

Along with changes in the inflammatory response, we found that although uptake of conidia into the cell was not impaired in CF macrophages, fungal killing was. Killing defects have been identified previously in CF macrophages in response to bacterial pathogens; however, CF monocytes and neutrophils from blood have been shown to kill fungi as efficiently as healthy controls ^31,51,52^. We have previously shown that failure to kill *Aspergillus* at the conidial stage leads inexorably to fungal germination within the cell resulting in inflammatory necroptosis and that this process is NFAT-dependent ^45^. We observed that *Aspergillus* infection induced more necrosis in CF macrophages than in wild-type equivalents. Necroptosis is regulated by the calcineurin-NFAT pathway with inhibition of this pathway by the calcineurin inhibitor FK506 (Tacrolimus) blocking necroptotic cell death ^45^. Our results suggest that in CF macrophages, dysregulated innate signalling leads to excessive programmed necrosis and a subsequent failure of control of fungal growth. These observations are consistent with previous studies, where blockade of IL1β signalling downstream of the inflammasome with Anakinra ameliorated both inflammation and fungal burden ^55^. Increased programmed necrosis in macrophages may be deleterious to the host, not only because of the associated loss of control of fungal outgrowth but also as lytic cell death releases inflammatory intra-cellular contents into the airway. Additionally, previous studies have highlighted a potential therapeutic role for Tacrolimus in the treatment of both fibrotic and allergic lung diseases. This, together with Tacrolimus’ inherent antifungal properties, suggest that targeted inhaled Tacrolimus delivery to the lung presents a novel area to investigate drug repurposing for treatment of the hyperinflammatory and ABPA phenotypes seen in CF ^66–68^.

CFTR modulators are a life changing therapy for the majority of pwCF helping to improve lung function. These modulators have previously been shown to dampen *Aspergillus*- induced inflammation in CF cells by reducing ROS production, known to be increased in CF phagocytes, without impairing fungal killing *in vitro* ^52,69^. Therefore, modulators may not only restore normal CFTR function in epithelial cells but also act as anti-inflammatory agents. We did not assess CFTR modulators in our murine infection model as despite showing a CF hyperinflammatory phenotype ^55,70,71^, in the context of this being a CFTR knockout, it lacks murine CFTR for the modulators to target. Of note, it has been shown that these animals may express human CFTR in both bone-marrow derived and alveolar macrophages under certain conditions due to the gut correction of the model required for survival ^72,73,39^. Therefore, future studies assessing the impact of CFTR modulators on the hyperinflammatory immune response to infection should focus on CF mouse strains in which, for example, the human F508del mutation has been modelled. ^72,73,39,74–76^.

In conclusion, the results of our *in vivo* studies show that loss of normal CFTR function increases the inflammatory response in the lung, in both acute and allergic aspergillosis models. *In vitro* data from both human and murine CF macrophages showed increased levels of NFκB and NFAT transcription factors and release of inflammatory cytokines. These increased levels of inflammation were associated with poor fungal control and increased levels of lytic cell death. Together these data suggests that CFTR dysfunction results in increased inflammatory signalling, with reduced antimicrobial control facilitating the lytic cell death of phagocytes. Given the observation that “pan-optosis” occurs in response to *A. fumigatus* ^77,78,46^, further exploration of therapeutic targeting of these pathways to limit lytic cell death may provide novel ways to reduce the inflammatory phenotype seen in CF in response to infection.

